# Insights into the evolution of symbiosis gene copy number and distribution from a chromosome-scale *Lotus japonicus* Gifu genome sequence

**DOI:** 10.1101/2020.04.17.042473

**Authors:** Nadia Kamal, Terry Mun, Dugald Reid, Jie-shun Lin, Turgut Yigit Akyol, Niels Sandal, Torben Asp, Hideki Hirakawa, Jens Stougaard, Klaus F. X. Mayer, Shusei Sato, Stig Uggerhøj Andersen

## Abstract

**Aim:** *Lotus japonicus* is a herbaceous perennial legume that has been used extensively as a genetically tractable model system for deciphering the molecular genetics of symbiotic nitrogen fixation. Our aim is to improve the *L. japonicus* reference genome sequence, which has so far been based on Sanger and Illumina sequencing reads from the *L. japonicus* accession MG-20 and contained a large fraction of unanchored contigs.

**Methods and Results:** Here, we use long PacBio reads from *L. japonicus* Gifu combined with Hi-C data and new high-density genetic maps to generate a high-quality chromosome-scale reference genome assembly for *L. japonicus*. The assembly comprises 554 megabases of which 549 were assigned to six pseudomolecules that appear complete with telomeric repeats at their extremes and large centromeric regions with low gene density.

**Conclusion and Perspectives:** The new *L. japonicus* Gifu reference genome and associated expression data represent valuable resources for legume functional and comparative genomics. Here, we provide a first example by showing that the symbiotic islands recently described in *Medicago truncatula* do not appear to be conserved in *L. japonicus*.

## Introduction

The roots of most plants are colonized by mycorrhizal fungi. This symbiotic interaction is ancient, perhaps dating back to the origin of land plants, and many of its genetic components have been co-opted to allow symbiotic nitrogen fixation in legumes ^1^. Much of the overlapping genetic framework, as well as components specific to both types of symbioses, have been uncovered using the model legumes *Lotus japonicus* (Lotus) and *Medicago truncatula* (Medicago) ^2^. Lotus is a perennial legume that has a short generation time, abundant flowers, and a small diploid genome with an estimated size of ~500Mb ^3^. In addition, Lotus is self-compatible and amenable to tissue culture and *Agrobacterium* transformation ^4^. It has been used very successfully for forward genetic studies, resulting in the first identification of a plant gene (*Nin*) required for nodulation ^5^, and the discovery of receptors for rhizobium Nod factors (NFR1 and NFR5) ^6^ and exopolysaccharides (EPR3) ^7^.

Lotus is also interesting from a legume phylogenetic point of view, as it is a member of the Robinoid clade, which lacks other species with comprehensive genetic and genomic resources. The Robinoids are part of the larger Hologalegina clade, which also includes the IRLC clade that comprises Medicago and important crops such as pea (*Pisum sativum*), chickpea (*Cicer arietinum*), alfalfa (*Medicago sativa*), and white clover (*Trifolium repens*) ^8^. The Hologalegina clade is sister to the Indigoferoid/Milettioid clade that includes soybean (*Glycine max*), common bean (*Phaseolus vulgaris*), pigeon pea (*Cajanus cajan*) and cowpea (*Vigna unguiculata*) ^8^. All these species engage in symbiotic nitrogen fixation, but their root nodule morphology differs. The Indigoferoid/Milettioid species soybean and common bean and the Robinoid species Lotus produce round, determinate nodules, while the IRLC legumes instead form elongated, indeterminate nodules with persistent meristems ^9^. High quality genetic and genomic Lotus resources will thus nicely complement those of other well-characterised legume species, facilitating functional, comparative and phylo-genomic studies of symbiotic nitrogen fixation, arbuscular mycorrhization and other legume traits of interest.

The genetic resources already available for Lotus include sequenced natural accessions ^10^ and recombinant inbred lines ^11,12^, as well as extensive populations of TILLING lines ^13^ and *LORE1* insertion mutants ^14^ In addition, large volumes of Lotus expression and *LORE1* data have been integrated in the online portal *Lotus* Base ^15^ (https://lotus.au.dk). Two Lotus accessions, MG-20 and Gifu B-129 (Gifu), have been especially frequently used ^16^. So far, genome sequencing efforts have focused exclusively on MG-20, resulting in the release of version 1.0, 2.5 and 3.0 MG-20 assemblies ^17^ (https://www.kazusa.or.jp/lotus/ and https://lotus.au.dk/). MG-20 v.3.0 is a hybrid assembly based on Sanger and Illumina data that comprises 132 scaffolds covering 232 Mbp aligned to the six Lotus chromosomes and an additional 162 Mbp of sequence in 23,572 unanchored contigs. This MG-20 assembly has proved very useful for genetic mapping and for genome-wide transcriptome, methylation and insertion mutant analyses ^7^,^14,18,19^, but it remains incomplete. Gifu originates from central Japan and is closely related to most of the sequenced accessions ^10^, whereas MG-20 is an atypical Lotus accession that originates from Miyakojima Island in the far south of Japan close to Taiwan. Considering also that the *LORE1* insertion mutant collection ^14^ was generated in the Gifu background, a high quality Lotus Gifu reference genome would not only facilitate comparative genomics studies, but also serve to underpin improvement of functional genomics and intraspecific diversity resources in Lotus.

Here, we present a high-quality Lotus Gifu reference assembly constructed based on ~100x PacBio read coverage and scaffolded using Hi-C and high-resolution genetic map data. We use this high-quality assembly to explore the positional clustering of putative orthologs of Medicago lncRNAs and compare nodule-regulated gene clusters between Lotus and Medicago. Conserved gene regulation was found for root and nodule samples, but evidence supporting conservation of the symbiotic islands discovered in Medicago did not emerge.

## Materials and Methods

### PacBio data generation and assembly

Lotus Gifu high-molecular weight DNA was extracted as described ^20^ and sent to Earlham Institute and Takara Bio Inc. for PacBio sequencing. A total of 11.8 million reads with an average length of 8 kb were generated. The PacBio reads were assembled using Canu (v1.3) ^21^ with the parameters corOutCoverage=100 errorRate=0.015 corMhapSensitivity=normal corMaxEvidenceErate=0.15 oeaMemory=15 cnsMemory=40. The assembled contigs were then polished using PacificBiosciences’ GenomicConsensus package using Quiver (https://github.com/PacificBiosciences/GenomicConsensus).

### Constructing genetic maps based on data from two RIL populations

Paired-end reads from recombinant inbred lines (RILs) of Gifu×*Lotus burttii* and Gifu×MG-20, as well as those from their respective parental lines (Lotus Gifu, Lotus MG-20, and *L. burttii*), were mapped to the polished assembly using BWA-MEM ^22^. Picard (http://broadinstitute.github.io/picard/) was used to dedupe the generated BAM files, followed by variant calling using mpileup provided by SAMtools ^23^. The resulting VCF files were filtered based on the following criteria: (1) minimum quality of 30, (2) minimum depth of 50, (3) must be biallelic, and (4) cannot contain missing genotypes. To improve the quality of the genetic map, further filtering was performed using a Python script to select solely for single nucleotide polymorphisms (SNPs) that are homozygous in the Gifu parent and homozygous alternative in the second RIL parent (MG-20 or *L. burttii*). To generate a consensus genotype call pattern for each contig across each RIL population (Gifu × *L. burttii* and Gifu×MG-20), the most commonly occurring genotype across all positions was selected.

### Assembly scaffolding based on genetic maps and Hi-C data

Gifu leaf tissue was sent to Phase Genomics (https://phasegenomics.com), where Hi-C sequencing was carried out and a draft proximity-based (Proximo) scaffolding generated. Chromatin conformation capture data was generated using a Phase Genomics (Seattle, WA) Proximo Hi-C ^24^. Intact cells from two samples were crosslinked using a formaldehyde solution, digested using the *Sau3AI* restriction enzyme, and proximity ligated with biotinylated nucleotides to create chimeric molecules composed of fragments from different regions of the genome that were physically proximal in vivo, but not necessarily genomically proximal. Molecules were pulled down with streptavidin beads and processed into an Illumina-compatible sequencing library. Sequencing was performed on an Illumina NextSeq 500, generating a total of 175,495,827 PE150 read pairs. Reads were aligned to the draft PacBio assembly scaffoldSeq.fasta using bwa mem with the −5 option ^22^. Alignments were then filtered with SAMtools ^23^ using the −F 2316 filtering flag.

Phase Genomics’ Proximo Hi-C genome scaffolding platform was used to create chromosome-scale scaffolds from the draft assembly in a method similar to that described by Bickhart et al. ^25^. As in the LACHESIS method ^26^, this process computes a contact frequency matrix from the aligned Hi-C read pairs, normalized by the number of *Sau3AI* restriction sites (GATC) on each contig, and constructs scaffolds in such a way as to optimize expected contact frequency and other statistical patterns in Hi-C data. Approximately 88,000 separate Proximo runs were performed to optimize the number of scaffolds and scaffold construction in order to make the scaffolds as concordant with the observed Hi-C data as possible. This process resulted in a set of six chromosome-scale scaffolds containing 549 Mbp of sequence (>99% of the draft assembly). Chimeric contigs were identified based on genetic map, Hi-C, and PacBio coverage data and split. The initial scaffolding was then iteratively improved using genetic map data followed by re-running Proximo scaffolding until genetic map and proximity-based scaffolding results converged.

### Genome annotation

The annotation of the Lotus Gifu genome was performed using evidence from transcriptome data as well as homology information from related species. For the homology-based annotation, available *Arabidopsis thaliana* (Araport11), *Glycine max* (v2.1) and Medicago (MtrunA17r5.0-ANR) protein sequences were combined. These protein sequences were mapped to the Lotus Gifu reference genome sequence using the splice-aware alignment tool GenomeThreader ^27^ (version 1.6.6; with the arguments -startcodon -finalstopcodon -species rice -gcmincoverage 70 -prseedlength 7 -prhdist 4). In the expression data-based step, multiple RNA-seq datasets (SRP127678, SRP105404, DRP000629, PRJNA622801) were used as evidence for the genome-guided prediction of gene structures. Therefore, reads from RNA-seq datasets were mapped to the genome using Hisat2 (version 2.1, parameter –dta) ^28^ and subsequently assembled into transcript sequences with Stringtie (version 1.2.3, parameters - m 150 -t -f 0.3) ^29^. Next, Transdecoder (version 3.0.0) (https://github.com/TransDecoder/TransDecoder) was used to identify potential open reading frames and predict protein sequences. Using BLASTP (ncbi-blast-2.3.0+, parameters -max_target_seqs 1 -evalue 1e-05) ^30^ the predicted protein sequences were compared against a protein reference database (UniProt Magnoliophyta, reviewed/Swiss-Prot) and used hmmscan (version 3.1b2) ^31^ to identify conserved protein family domains for all proteins. BLAST and hmmscan results were then used by Transdecoder-predict and the best translations per transcript sequence was selected. Finally, results from the two gene prediction approaches were combined and redundant protein sequences were removed. Additionally, some symbiosis genes were manually curated (**Supplemental table 6**).

In order to classify gene models into complete and functional genes, non-coding transcripts, pseudogenes and transposable elements, a confidence classification protocol was applied. Candidate protein sequences were compared against the following three databases using BLAST: PTREP, a manually curated database of hypothetical proteins that contains deduced protein sequences, from which frameshifts have mostly been removed (http://botserv2.uzh.ch/kelldata/trep-db/index.html); a database with annotated proteins from the legumes *Glycine max* and Medicago, called ‘Fab’ hereafter; and UniMag, a database of validated proteins from the Magnoliophyta. UniMag protein sequences were downloaded from UniProt and further filtered for complete sequences with start and stop codons. Best hits were selected for each predicted protein to each of the three databases. Only hits with an E-value below 10e-10 were considered. Furthermore, only hits with subject coverage above 80% were considered significant and protein sequences were further classified into high and low confidence. High confidence (HC) protein sequences are complete and have a subject and query coverage above the threshold in the UniMag database (HC1) or no blast hit in UniMag but in Fab and not PTREP (HC2). While a low confidence (LC) protein sequence is not complete and has a hit in the UniMag or Fab database but not in PTREP (LC1), or no hit in UniMag and Fab and PTREP but the protein sequence is complete. Functional annotation of transcripts as well as the assignment of GO terms was performed using the tool “Automatic assignment of Human Readable Descriptions - AHRD”. AHRD performs BLASTP search against Swiss-Prot, The Arabidopsis Information Resource (TAIR) and TrEMBL databases to perform functional annotation based on homology to other known proteins and integrates domain search results from InterProScan as well as gene ontology (GO) terms ^32^. Repeats were annotated using RepeatMasker ^33^ version 3.3 with a custom Fabaceae-library in sensitive mode. Non-coding RNAs were predicted using tRNAscan-SE (version 1.3.1) ^34^, RNAmmer (version 1.2) ^35^ and Infernal (version 1.1.2) ^36^ with default parameters. The results were merged subsequently.

### Expression atlas

Raw Lotus Gifu RNA-seq reads were obtained from either the Sequence Read Archive (SRA) for the listed accessions or generated in this study (**Supplemental table 1**). For data in this study, three day old Lotus Gifu seedlings were transferred to filter paper covered agar (1.4% agar noble) slants. Roots were treated with *M. loti* R7A, 6-Benzylaminopurine (1 μM) or mock and a 1 cm segment of root tissue corresponding to the zone of emerging root hairs at time of treatment was harvested. For nodule tissue, whole nodules were harvested. Libraries were constructed and sequenced by Novogene (Hong Kong) using PE-150bp reads on the Illumina NovaSeq 6000 instrument. A decoy-aware index was built for Gifu transcripts using default Salmon parameters and reads were quantified using the --validateMappings flag ^37^ (Salmon version 0.14.1). A normalised expression atlas across all conditions was constructed using the R-package DESeq2 version 1.20 ^38^ after summarising gene level abundance using the R-package tximport (version 1.8.0). Normalised count data obtained from DESeq2 are available in the *Lotus* Base expression atlas (https://lotus.au.dk/expat/) ^15^.

### Analysis of symbiotic islands

Medicago A17 proteins associated with symbiotic islands as defined by Pecrix et al. ^39^, were blasted against Lotus Gifu proteins annotated in the present assembly, and the best hit was extracted. It was then determined if there was microsynteny between the Medicago A17 genes in the symbiotic island and the best Lotus Gifu matches (**Supplemental file 5**). Medicago A17 RNA-seq data (**Supplemental table 2**) was trimmed using trimmomatic (10.1093/bioinformatics/btu170), trimmed reads were mapped to the Medicago A17 v.5 reference sequence (MtrunA17r5.0) using the splice aware STAR aligner (version 2.5.1a) ^40^. A read was allowed to map in at most 10 locations (–outFilterMultimapNmax 10) with a maximum of 4% mismatches (–outFilterMismatchNoverLmax 0.04) and all non--canonical intron motifs were filtered out (–outFilterIntronMotifs RemoveNoncanonicalUnannotated). In order to obtain non-unique gene-level counts from the mapping files, HTSeq (version 0.9.1) ^41^ with the ‘nonunique all’-method was used. Normalization of read counts was performed by library sequence depth using the R-package DESeq2 (version 1.23.3) ^38^.

Log expression ratios of 10 days post inoculation (dpi) nodule samples versus non-inoculated root samples were calculated for Lotus and Medicago and Pearson correlation coefficients were calculated (**Supplemental tables 1–2**). For calculation of Pearson correlation coefficients, all Medicago A17 RNA-seq samples listed in **Supplemental table 2** were used, while only Lotus Gifu root and nodule samples were used (**Supplemental table 1**). When analysing the largest possible set of genes (**Figure 3B**), all Medicago A17 genes with a match to a Lotus Gifu gene anywhere in the genome were included along with one Lotus Gifu match per Medicago A17 gene, allowing many copies of the same Lotus gene. For analysis of unique Lotus genes, only a single Medicago A17 gene was included per Lotus Gifu match within the microsyntenic region and islands with less than three Lotus Gifu microsyntenic hits were not considered (**Figure 3C**). All statistical analyses were carried out using R v. 3.4.3. The scripts used for analysis are freely available from GitHub (https://github.com/stiguandersen/LotjaGifuGenome).

### Data availability

Sequencing data is available from SRA. PacBio data used for genome assembly and Hi-C data from Phase Genomics used for construction of proximity map (PRJNA498060); Illumina paired-end data from RIL resequencing used for genetic map construction (PRJNA498068); *L. burttii* genomic DNA reads (PRJNA635235); RNA-seq data used for annotation (PRJNA622801); RNA-seq expression atlas data (PRJNA622396). Assembly pseudomolecules are available from the NCBI Nucleotide repository with accession numbers AP022629-AP022637. Pseudomolecule sequences and genome annotation information are also found in Supplemental Files 2 and 3 and are available for browsing and download at *Lotus* Base (https://lotus.au.dk) and LegumeBase (https://www.legumebase.brc.miyazaki-u.ac.jp) and for synteny comparisons at CoGe (https://genomevolution.org/coge/GenomeInfo.pl?gid=58121).

## Results and data description

### A chromosome-scale Lotus Gifu assembly including telo- and centromeric repeats

We generated a total of 11.8 million PacBio RSII reads, which we assembled using Canu ^21^ into 1,686 contigs with an N50 of 807 kb and a total length of 554 Mb (**Table 1**). We first scaffolded the contigs using 175 million Proximo Hi-C reads (Phase genomics). To validate the scaffolding, we mapped whole genome re-sequencing data from two recombinant inbred line populations ^12^ to the PacBio contigs. The vast majority of the assembly, 99.5%, was contained within contigs that had at least one polymorphic SNP marker, leaving only 2.5 Mb of sequence on markerless contigs (**Table 1**). We compared the Hi-C scaffolding results to the genetic maps generated based on the recombinant inbred line data (Supplemental file 1) and moved contigs according to genetic linkage. We then repeated the scaffolding until the Proximo Hi-C results were concordant with the genetic maps and the contigs were arranged in six pseudomolecules corresponding to the six Lotus chromosomes (Supplemental file 2). The total length of the assembly was close to the expected genome size of ~500 Mb (**Table 1**), and we found canonical telomeric repeats at the ends of all pseudomolecules, except for the bottom of chromosome three, indicating a high completeness of the assembly. The 2.5 Mb of unanchored contigs placed on chr0 contained a substantial amount of pericentromeric repeats.

**Table 1.**
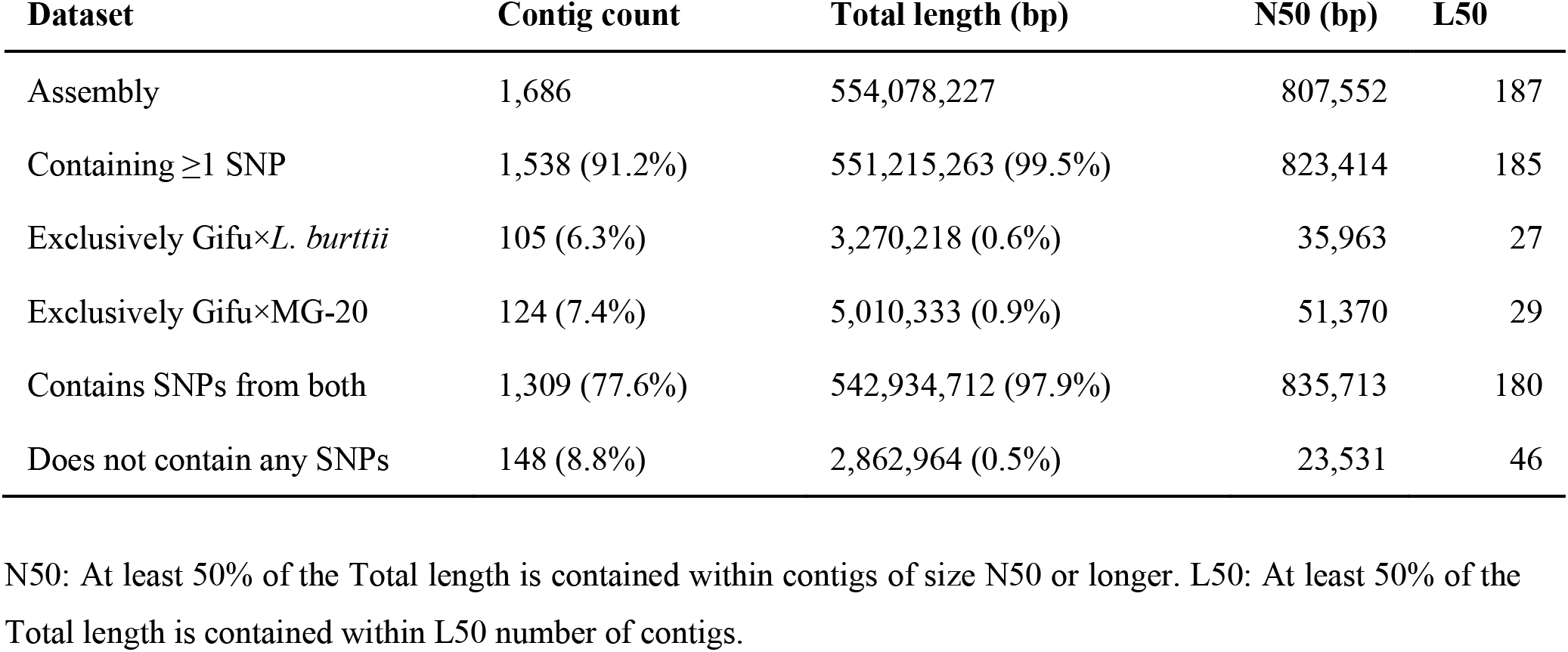
Assembly and genetic map statistics

| Dataset | Contig count | Total length (bp) | N50 (bp) | L50 |
| --- | --- | --- | --- | --- |
| Assembly | 1,686 | 554,078,227 | 807,552 | 187 |
| Containing $\geq 1$ SNP | 1,538 (91.2%) | 551,215,263 (99.5%) | 823,414 | 185 |
| Exclusively Gifu×*L. burttii* | 105 (6.3%) | 3,270,218 (0.6%) | 35,963 | 27 |
| Exclusively Gifu×MG-20 | 124 (7.4%) | 5,010,333 (0.9%) | 51,370 | 29 |
| Contains SNPs from both | 1,309 (77.6%) | 542,934,712 (97.9%) | 835,713 | 180 |
| Does not contain any SNPs | 148 (8.8%) | 2,862,964 (0.5%) | 23,531 | 46 |

N50: At least 50% of the Total length is contained within contigs of size N50 or longer. L50: At least 50% of the Total length is contained within L50 number of contigs.

Regarding the highly repetitive sequences, three 45S rDNA clusters and a 5S rDNA gene cluster were anchored on chromosomes 2, 5 and 6, and on chromosome 2, respectively, consistent with FISH data (**Figure 1A**) ^42^. In addition to the regions with a high density of repetitive sequences, corresponding to the pericentromeric regions of each chromosome, small regions with high densities of repetitive sequences were identified within the gene rich regions at the bottom arm of chromosomes 2 and 4 (**Figure 1A**). The location of these regions corresponded to the positions of chromosome knobs reported in the previous cytological analyses ^42,43^. These regions with highly dense repetitive sequences tend to be composed of contigs with short length, and thus a significant number of the sequence gaps (389 out of 1555) were found in these regions. Despite the relatively high frequency of sequence gaps in these repetitive regions, the Hi-C reads provided sufficient physical linking information to allow scaffolding.

**Figure 1:**
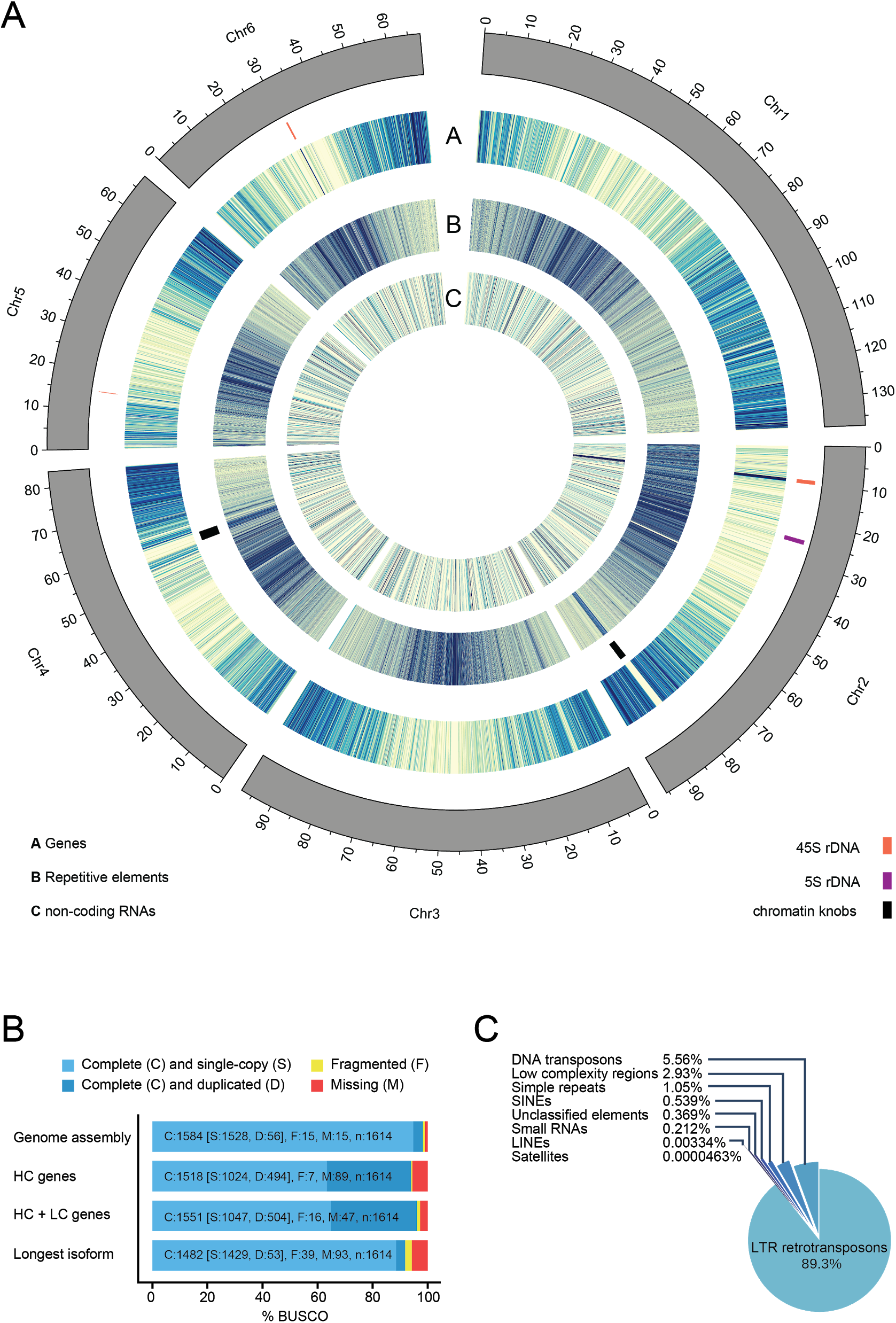
**A**) Circos diagram displaying heatmaps of the numbers of genes and ncRNAs (100 Mb bins) and bases covered by repetitive elements (10 Mb bins) in the Lotus Gifu genome. **B**) BUSCO v4 scores of the Lotus Gifu assembly (98.2%), the high confidence gene set (94%), the high- and low confidence gene set (96.1%) and of only the longest transcript of each gene (91.8%) from the joint high and low confidence gene set. Lineage used: embryophyta_odb10. **C**) Distribution of repetitive elements in the Lotus Gifu genome.

### Genome annotation

Based on evidence from expression data as well as homology information from related species, 30,243 genes were annotated, 21,778 of which represent high confidence gene models (**Table 2**, **Supplemental file 3**). Using the embryophyta_odb10 lineage 1,584 out of 1,614 (98.2%) complete BUSCO v4 orthologs ^44^ were found in the genome assembly and 1,551 (96.1%), were identified within the annotated gene set (**Figure 1B**). The high confidence gene set had a BUSCO score of 94%. Using AHRD ^32^ we could assign functional annotations to 29,429 genes (97%). Of these, 70.53% fulfilled all three AHRD quality criteria, 16.85% fulfilled two and 11.8% fulfilled one criterion. We then annotated non-coding RNAs, identifying 2,933 in total that comprised 128 micro RNAs, 851 snoRNAs, 88 tRNAs, 795 rRNAs and others. In total, gene models covered 156,379,918 bases and coding exons covered 60,649,299 bases of the genome assembly.

**Table 2:** Genome annotation statistics

|  | **Lotus Gifu**<br>**v.1.2**<br>**HC+LC** | **Lotus Gifu**<br>**v.1.2 HC** | **Medicago**<br>**A17 v.4** | **Medicago**<br>**A17 v.4 HC** | **Medicago**<br>**A17 v.5** | ***Glycine max***<br>**Williams 82**<br>**v. 2.1** |
| --- | --- | --- | --- | --- | --- | --- |
| Number of genes | 30,243 | 21,778 | 50,444 | 31,451 | 51,316 | 52,872 |
| Number of coding genes | 29,554 | 21,778 | 50,444 | 31,451 | 44,623 | 52,872 |
| Number of mRNAs | 49,868 | 37,994 | 57,585 | 38,175 | 44,623 | 86,256 |
| Number of exons | 306,545 | 264,198 | 267,394 | 397,385 | 189,379 | 560,910 |
| Number of CDSs | 262,442 | 236,845 | 257,792 | 376,276 | 174,461 | 516,059 |
| Average CDS lengths (bp) | 1,216.2 | 1,385.4 | 1,038.4 | 1,272.4 | 1,017.7 | 1,350.7 |
| Average exon lengths (bp) | 417.54 | 373.77 | 282.58 | 261.92 | 360.25 | 312.48 |
| Average intron lengths (bp) | 527.12 | 513.71 | 444.41 | 438.41 | 476.57 | 519.19 |
| Average transcripts per gene | 1.65 | 1.74 | 1.14 | 1.54 | 1 | 1.63 |
| Average exons per transcript | 6.15 | 6.95 | 4.64 | 6.75 | 4.19 | 6.5 |
| Average CDS exons per transcript | 5.23 | 6.23 | 4.48 | 6.39 | 3.91 | 5.98 |

HC: high confidence gene models. LC: low confidence gene models.

Repetitive elements made up 260,312,827 bases (46.96%) of the genome. Of these, long terminal repeat retrotransposons accounted for most of the repeat content of the genome (42.51%), followed by DNA transposons and low complexity regions (**Figure 1C**). Chromosomes 1, 3, 4, 5, and 6 showed centrally located pericentromeric regions rich in repetitive elements flanked by gene-rich regions (**Figure 1A**). In contrast, the centromere of chromosome 2 appeared to be distally located near the top of the chromosome, which also carried a large cluster of rRNA genes (**Figure 1A**).

### RNA-seq based expression atlas

To produce a gene expression atlas, publicly available and new RNA-seq data from Lotus Gifu was obtained for 35 conditions across different tissues, symbiotic and pathogenic interactions (**Supplemental table 1**). The conditions available include root hair, nodule primordia and nodules obtained after inoculation with *Mesorhizobium loti* R7A and root interactions with microbes across a symbiont-pathogen spectrum ^18^; root and shoot tissues three days after roots were inoculated with *M. loti* ^45^; root symbiotic susceptible zone treated with cytokinin (1 μM 6-Benzylaminopurine (BA)) or *M. loti* R7A (this study); roots inoculated with the arbuscular mycorrhizal fungus (AMF), *Glomus intraradices* ^46^; root, leaf, immature flowers, mature flowers, pods and seeds (National Institute for Basic Biology, 2016). Gene-level quantification of the data was normalised across conditions (**Supplemental file 4**) and is made available through *Lotus* Base (https://lotus.au.dk/expat/) to provide a readily accessible expression viewer. Well-described nodulation genes showed the expected expression patterns across the conditions represented in the expression atlas (**Figure 2**).

**Figure 2.**
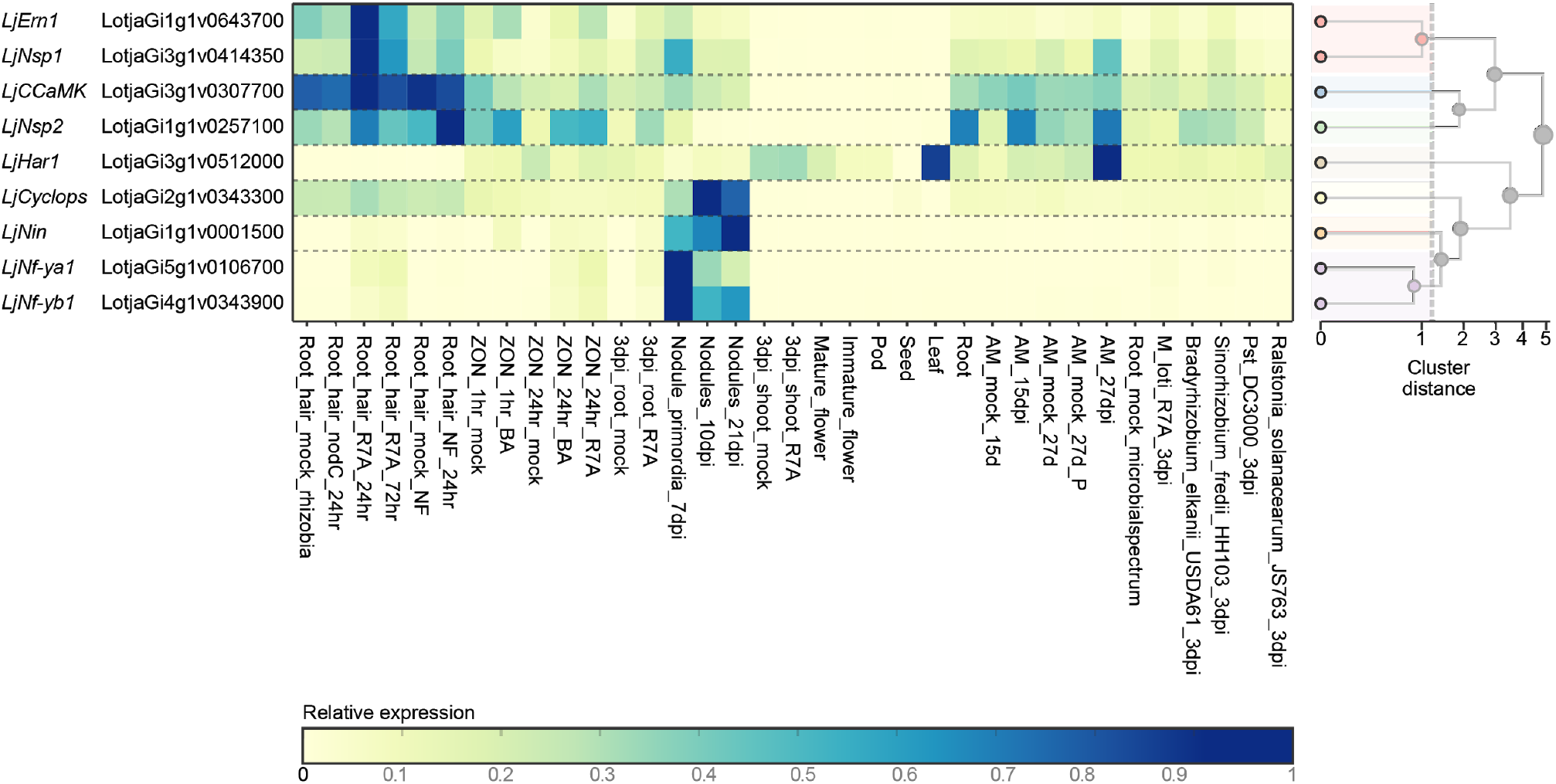
Expression profiles of known symbiosis genes. Expression values from the Lotus Gifu RNA-seq expression atlas are shown for the indicated genes. A full description of the conditions included is shown in **Supplemental table 1**. The heatmap was generated from https://lotus.au.dk/expat/ using the normalise by row function.

### Symbiotic islands are not generally conserved between Lotus and Medicago

Recently, “symbiotic islands” representing clusters of genes that showed co-regulated, symbiosis-related expression profiles were identified in Medicago A17 ^39^. Interestingly, these clusters were rich in long non-coding (lnc) RNAs, and it was proposed that the lncRNAs may be involved in regulating symbiosis-related gene expression. To investigate if the Medicago symbiotic islands were conserved in Lotus, we extracted the best Lotus Gifu BLAST hits against the Medicago A17 genes reported to reside within symbiotic islands (**Supplemental file 5**). Protein coding genes were generally well conserved and showed high levels of microsynteny, regardless of whether or not they were present in gene islands that showed symbiosis-related differential expression (**Table 3**). Out of 760 islands, 266 had at least three distinct Lotus Gifu hits in microsyntenic regions, and the region with the largest overlap comprised 12 hits. In contrast, most Medicago A17 lncRNAs had no putative orthologs in the Lotus Gifu genome, and, when identified, they were often not found within the designated microsyntenic region (**Table 3**). Across all 760 investigated islands, a total of six had two lncRNA hits to the Lotus Gifu microsyntenic region, and no island had more than two.

**Table 3.**
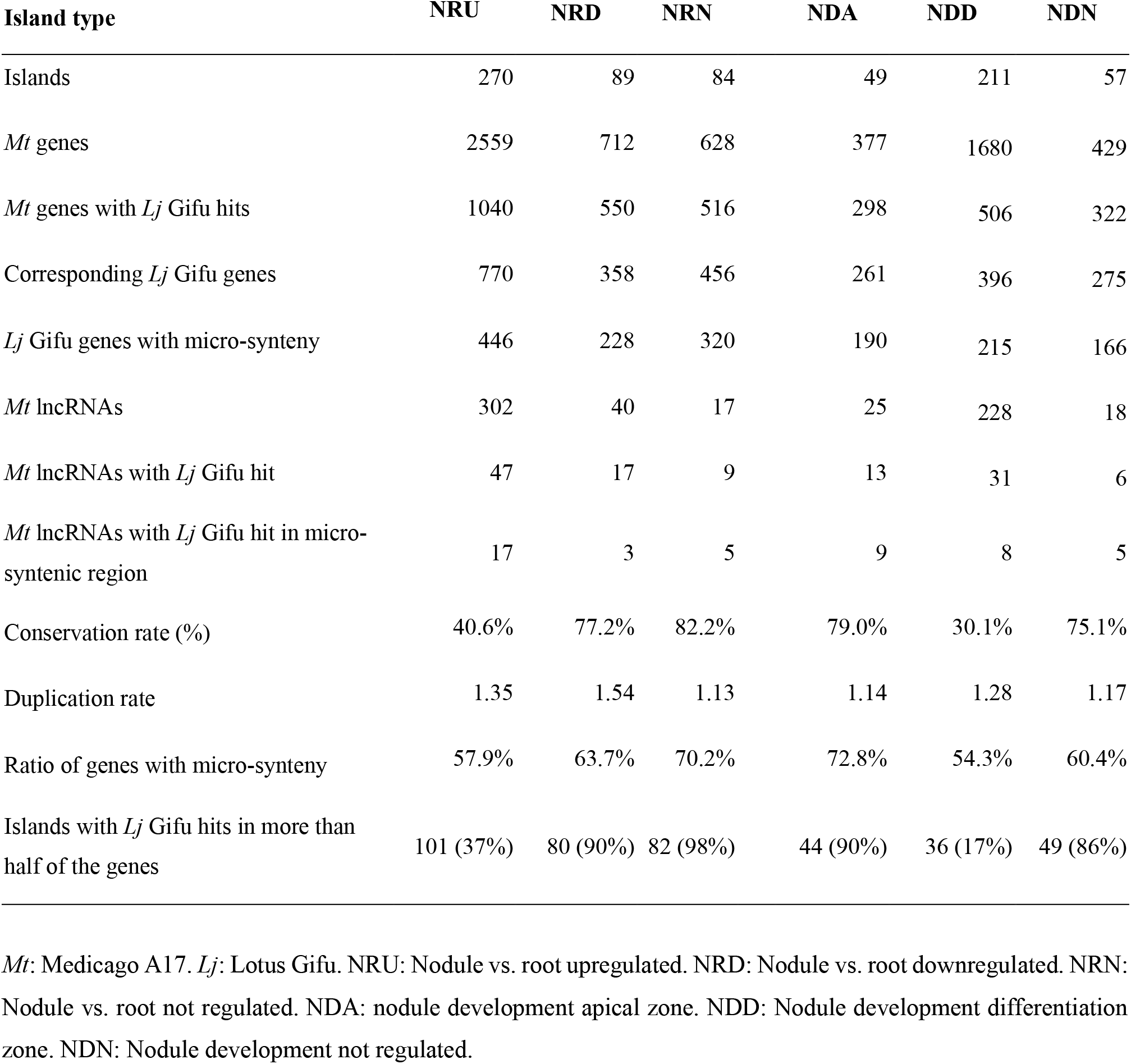
Conservation of symbiotic islands between Lotus and Medicago.

The limited conservation and lack of positional clustering make it unlikely that putative orthologs of Medicago lncRNAs are generally part of symbiotic islands in Lotus. Instead, we looked further into the protein coding genes to determine if their organisation into symbiotic islands could be conserved. All 760 islands contain at least one protein coding gene. Out of these, we examined the 443 islands associated with nodule-regulated genes designated “Nodule upregulated (NRU)”, “Nodule downregulated (NRD)” and “Nodule non-regulated (NRN)”. First, we investigated the level of expression conservation by comparing the expression of Medicago genes in symbiotic islands and their Lotus syntenic homologs in root and 10 dpi nodule samples (**Supplemental tables 1-2 and Supplemental files 4 and 6**). The genes associated with Medicago NRU islands showed strongly correlated expression responses in Lotus and Medicago, NRD genes showed a less pronounced correlation, while there was no correlation for the NRN genes (**Figure 3A**).

**Figure 3:**
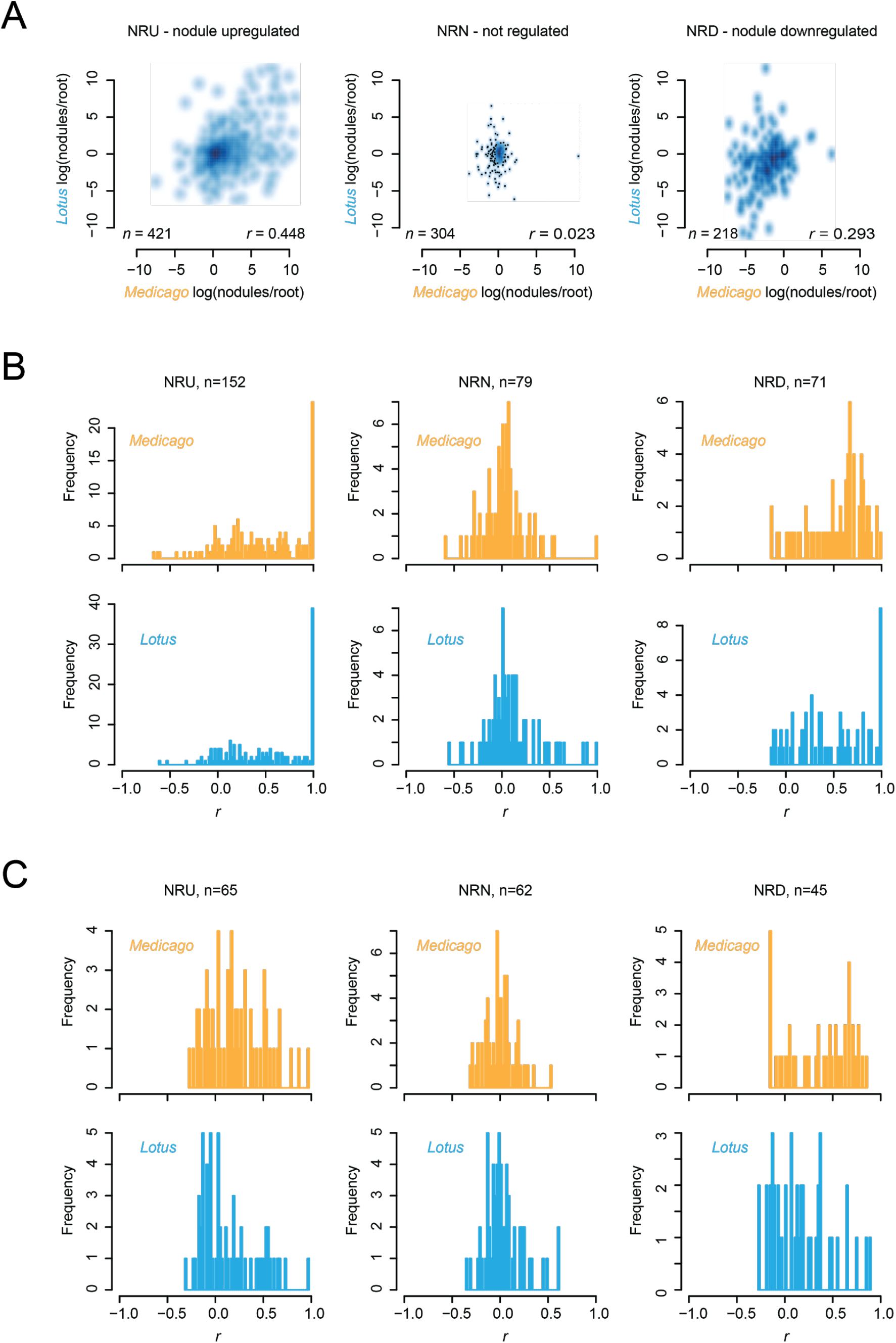
Symbiotic island gene expression. **A)** Log(nodule/root) expression ratios for genes in Medicago symbiotic islands and their best *Lotus* blast matches. *n*: gene count. *r*: Pearson correlation coefficient for the Lotus and Medicago log(nodule/root) ratios. **B-C)** Histograms of Pearson correlation coefficients for symbiotic islands. The Pearson correlation coefficient for each island is an average of the coefficients resulting from pairwise comparisons of the gene expression profiles of all genes residing within that island. *n*: symbiotic island count. **B)** All genes in Medicago symbiotic islands with a putative *Lotus* homolog with expression data. Multiple copies of the same *Lotus* gene are allowed. **C)** Only one *Lotus* copy and one corresponding Medicago gene is included in the analysis and it is further required that each *Lotus* island contains at least three members. *Lotus*: expression data from *Lotus*. Medicago: expression data from Medicago A17.

To quantify the level of co-regulation within putative symbiotic islands, we calculated the average Pearson correlation coefficients for each island based on the gene expression data from root and nodule samples (**Supplemental tables 1-2**). First, we included all genes in Medicago symbiotic islands that had a Lotus BLAST match anywhere in the genome along with their best Lotus match. If a Lotus gene was the best match for multiple Medicago genes, it was included multiple times in the analysis. Especially for the NRU islands, this resulted in a very pronounced skew towards high correlation coefficients as compared to the NRN islands, and this was true both for Lotus and Medicago (**Figure 3B**).

We then repeated the analysis including only Medicago-Lotus syntenic homolog pairs from islands with at least three unique Lotus genes. That is, if multiple Medicago genes matched the same Lotus gene, only a single Medicago gene was retained and each unique Lotus gene was only included once per island. This resulted in a marked reduction in the number of islands and the large peak of near perfect correlation coefficients for NRU islands disappeared for both Lotus and Medicago (**Figure 3C**). Since there was no longer a major difference between the root/nodule-based correlation coefficients between the nodule-regulated NRU and NRD islands and the NRN controls, it appears that local gene amplification in Medicago is a major cause of the symbiotic island signal. This is consistent with an overall high ratio of Medicago to Lotus genes in symbiotic islands (**Table 3**). Symbiotic islands are thus not generally conserved between Lotus and Medicago and are not general features of legume genomes. However, we did find a few examples of gene clusters that showed conserved co-regulation for root and nodule samples (**Supplemental tables 3-5**). In Lotus, NRU island SRI_NDD0105, which had the second highest Lotus correlation coefficient (**Supplemental table 3**), had three very similar copies of a nodulin gene, suggesting that local gene amplification also plays a role here. In contrast, the NRU island with the highest Lotus correlation coefficient (SRI_NRU0026) comprised three very different genes, perhaps warranting further investigation (**Supplemental table 3**).

## Conclusion and Future Perspectives

By applying long PacBio reads, the contiguity of the assembly was improved compared to the MG-20 version 3.0 assembly that was a hybrid assembly based on Sanger and Illumina sequences. Using Hi-C paired-end reads and high-density SNP marker information generated by re-sequencing of Lotus Gifu x *L. burttii* RILs, 1584 contigs were anchored onto 6 chromosomes with 42 scaffolds, providing a high-quality and well-validated assembly. The number of scaffolds was a bit larger than that of the latest Medicago A17 sequence (Mt5.0) due to manual correction of Hi-C scaffolding errors based on the SNP marker information. Typical Hi-C scaffolding errors were identified in the distal regions of each pseudomolecule and at the border regions of chromosome knobs located on chromosomes 2 and 4, presumably due to an atypical three-dimensional chromosome conformation in those regions. A total of 30,243 high and low confidence gene models were annotated, which corresponds approximately to the number of high confidence gene models in the Medicago v. 4 assembly (**Table 2**). The total number of annotated genes is higher for Medicago versions 4 and 5 than for the current Lotus Gifu assembly. However, the number of exons per transcript is markedly lower for the full Medicago gene sets than for the Lotus gene and Medicago v.4 high confidence gene sets, suggesting that the differences in gene numbers are due to different stringencies in including small genes with few exons. As expected, the paleopolyploid soybean (*Glycine max*) ^47^ has a higher number of annotated genes than Lotus but retains a similar exon per transcript ratio despite more than 50,000 annotated genes.

The availability of a high-quality Lotus Gifu assembly will facilitate further improvements of genetic and genomic Lotus resources. The *LORE1* mutant collection, which includes more than 130,000 insertion mutant lines, is in the Gifu genetic background, but was annotated based on the MG-20 v. 3.0 assembly ^14^. Using the new Gifu sequence, the *LORE1* insertions can now be more accurately characterised. Likewise, Gifu is more closely related to the majority of the collection of natural Lotus accessions that was recently characterized ^10^, and the new reference assembly should allow an improved characterization of the genetic diversity. Here, we have mapped existing and new RNA-seq data to the Gifu assembly to provide a consistently normalized and updated Lotus gene expression atlas readily available through *Lotus* Base ^15^. The current atlas does not comprise as many samples as previously profiled using microarrays ^48,49^, but it is not limited by probe set selection and includes data on all annotated and expressed genes.

The new assembly and expression atlas also proved useful in interspecific comparisons, since the complete pseudomolecules allowed us to accurately assess synteny with Medicago to investigate the level of conservation of plant symbiotic islands. Interestingly, the recently identified Medicago symbiotic islands did not appear to be conserved in Lotus. This was most evident for the Medicago non-coding RNAs, for which we could find only very few matching sequences in Lotus despite the completeness of the assembly. It should be noted that many of the transcripts classified as long non-coding RNAs in the Medicago study ^39^ in fact encode peptides, most notably the large family of nodule cysteine-rich (NCR) peptides. The NCR peptides are characteristic of the Inverted Repeat Lacking Clade (IRLC) legume lineage and thus not found in Lotus ^50^. The same appears to be the case for the other transcripts in the non-coding class, indicating that non-coding and peptide-encoding genes have evolved rapidly and are not generally required for legume-rhizobium symbiosis across determinate and indeterminate nodulators. For the protein coding genes in symbiotic islands, we found much higher levels of conservation and microsynteny, but most of the local co-regulation appeared to be related to tandem gene duplications in Medicago. Generally, Medicago seems to have experienced not only a rapid expansion of NCR peptide genes and lncRNAs involved in symbiosis, but also of protein coding genes with symbiosis-related expression patterns, and our results clearly indicate that symbiotic islands are not general features of legume genomes.

The analysis of symbiotic islands represents only a first use case for the new high-quality Lotus genomic data, and we anticipate that it will be broadly used in genomics studies. The data will included in comparative genomics websites such as Phytozome ^51^ and Legume Information System ^52^ and it is already available at CoGe (https://genomevolution.org/coge/GenomeInfo.pl?gid=58121) ^53^. In addition, the high completeness of the assembly and geneset makes the data well suited for phylogenomic studies that rely on precise genomic data for large-scale cross-species analyses ^54^.

## Author contributions

Conceptualization, S.U.A., S.S. and K.F.X.M; Validation, N.K., S.S., K.F.X.M and S.U.A.; Formal Analysis, N.K., T.M., D.R., T.Y.A., T.A., H.H., S.S. and SUA; Investigation, N.S; Resources, J.S., S.S., K.F.X.M., S.U.A.; Data Curation, N.K., S.S., K.F.X.M and S.U.A.; Writing – Original Draft, S.U.A. and N.K.; Writing – Review & Editing, S.U.A. and N.K. with input from all authors; Visualization, N.K., S.U.A. and D.R.; Supervision, S.U.A., S.S. and K.F.X.M.; Project Administration, J.S., S.S., K.F.X.M. and S.U.A.; Funding Acquisition, J.S., S.S., K.F.X.M., S.U.A.

## Acknowledgements

The work was supported by the Danish National Research Foundation grant DNRF79 (J.S.), the Genome Information Upgrading Program of the National BioResource Project in 2014 and 2015 (S.S.), a JST CREST grant (number JPMJCR16O1) (S.S.), the CRISBAR grant of the German ministry for education and research (BMBF) (K.F.X.M.) and grant no. 10-081677 from The Danish Council for Independent Research | Technology and Production Sciences (S.U.A.).

## Supplemental files

**Supplemental file 1**: Contig assignment to genetic map segments

**Supplemental file 2**: Lotus Gifu version 1.1 genome sequence

**Supplemental file 3**: Lotus Gifu version 1.2 genome annotation

**Supplemental file 4**: Lotus Gifu expression atlas

**Supplemental file 5**: Lotus Gifu BLAST matches to Medicago A17 genes in symbiotic islands

**Supplemental file 6**: Medicago A17 expression data

## Supplemental information

### Supplemental tables

**Supplemental table 1.** *Lotus* Gifu RNA-seq samples. Nod regulation: Samples used for calculating expression differences between roots and nodules. Correlation: Samples used for calculating Pearson correlation coefficients for gene expression co-regulation in symbiotic islands. dpi: days post inoculation. NIBB: National Institute for Basic Biology, Japan.

| Tissue | Sample name | Nod regulation | Correlation | Replicates | Reference | Bioproject ID |
| --- | --- | --- | --- | --- | --- | --- |
| Root susceptible zone | ZON_1hr_mock |  |  | 3 | Current study | PRJNA622396 |
| Root susceptible zone | ZON_1hr_BA |  |  | 3 | Current study | PRJNA622396 |
| Root susceptible zone | ZON_24hr_mock |  |  | 3 | Current study | PRJNA622396 |
| Root susceptible zone | ZON_2 hr_BA |  |  | 3 | Current study | PRJNA622396 |
| Root susceptible zone | ZON_24hr_R7A |  |  | 3 | Current study | PRJNA622396 |
| Nodule | nodules_10dpi | x | x | 3 | Current study | PRJNA622396 |
| Root | 3dpi_root_mock | x | x | 3 | Munch et al., 2018 | PRJNA384655 |
| Root | 3dpi_root_R7A |  | x | 3 | Munch et al., 2018 | PRJNA384655 |
| Shoot | 3dpi_shoot_mock |  |  | 3 | Munch et al., 2018 | PRJNA384655 |
| Shoot | 3dpi_shoot_R7A |  |  | 3 | Munch et al., 2018 | PRJNA384655 |
| Flower | Mature_flower |  |  | 3 | NIBB | PRJDB2436 |
| Flower | Immature_flower |  |  | 3 | NIBB | PRJDB2436 |
| Pod | Pod |  |  | 3 | NIBB | PRJDB2436 |
| Seed | seed |  |  | 3 | NIBB | PRJDB2436 |
| Root | root |  | x | 3 | NIBB | PRJDB2436 |
| Leaf | leaf |  |  | 3 | NIBB | PRJDB2436 |
| Root | AM_15dpi |  |  | 3 | Handa et al., 2015 | PRJDB2819, PRJDB2576, PRJDB3212 |
| Root | AM_27dpi |  |  | 3 | Handa et al., 2015 | PRJDB2819, PRJDB2576, PRJDB3212 |
| Root | AM_mock_15d |  |  | 3 | Handa et al., 2015 | PRJDB2819, PRJDB2576, PRJDB3212 |
| Root | AM_mock_27d_P |  |  | 3 | Handa et al., 2015 | PRJDB2819, PRJDB2576, PRJDB3212 |
| Root | AM_mock_27 |  |  | 3 | Handa et al., 2015 | PRJDB2819, PRJDB2576, PRJDB3212 |
| Root hair | root_hair_mock_rhizobia |  |  | 3 | Kelly et al., 2018 | PRJNA422278 |
| Root hair | root_hair_nodC_24h |  |  | 3 | Kelly et al., 2018 | PRJNA422278 |
| Root hair | root_hair_R7A_24hr |  |  | 3 | Kelly et al., 2018 | PRJNA422278 |
| Root hair | root_hair_R7A_72hr |  |  | 3 | Kelly et al., 2018 | PRJNA422278 |
| Root hair | root_hair_mock_NF |  |  | 3 | Kelly et al., 2018 | PRJNA422278 |
| Root hair | root_hair_NF_24hr |  |  | 3 | Kelly et al., 2018 | PRJNA422278 |
| Nodule | nodule_primordia_7_dpi |  | x | 3 | Kelly et al., 2018 | PRJNA422278 |
| Nodule | nodules_21dpi |  | x | 1 | Kelly et al., 2018 | PRJNA422278 |
| Root | Root_mock_microbialspectrum |  | x | 2 | Kelly et al., 2018 | PRJNA422278 |
| Root | M_loti_R7A_3dpi |  | x | 2 | Kelly et al., 2018 | PRJNA422278 |
| Root | Bradyrhizobium_elkanii_USDA61_3dpi |  |  | 2 | Kelly et al., 2018 | PRJNA422278 |
| Root | Sinorhizobium_fredii_HH103_3dpi |  |  | 2 | Kelly et al., 2018 | PRJNA422278 |
| Root | Pseudomonas_syringae_pv_tomato_DC3000_3dpi |  |  | 2 | Kelly et al., 2018 | PRJNA422278 |
| Root | Ralstonia_solanacearum_JS763_3dpi |  |  | 2 | Kelly et al., 2018 | PRJNA422278 |

**Supplemental table 2.** Medicago A17 RNA-seq data. SRA: Sequence read archive (https://www.ncbi.nlm.nih.gov/sra).

| SRA Sample ID | Sample Name | Tissue | Treatment |
| --- | --- | --- | --- |
| SRR5740859 | MtNod0dpi_1 | Roots | Non-inoculated |
| SRR5740858 | MtNod0dpi_2 | Roots | Non-inoculated |
| SRR5740868 | MtNod0dpi_3 | Roots | Non-inoculated |
| SRR5740875 | MtNod4dpi_1 | Roots | 4 days post inoculation with *S. meliloti* 1021 |
| SRR5740878 | MtNod4dpi_2 | Roots | 4 days post inoculation with *S. meliloti* 1021 |
| SRR5740877 | MtNod4dpi_3 | Roots | 4 days post inoculation with *S. meliloti* 1021 |
| SRR5740870 | MtNod10dpi_1 | Nodules | 10 days post inoculation with *S. meliloti* 1021 |
| SRR5740864 | MtNod10dpi_2 | Nodules | 10 days post inoculation with *S. meliloti* 1021 |
| SRR5740861 | MtNod10dpi_3 | Nodules | 10 days post inoculation with *S. meliloti* 1021 |
| SRR5740862 | MtNod14dpi_1 | Nodules | 14 days post inoculation with *S. meliloti* 1021 |
| SRR5740866 | MtNod14dpi_2 | Nodules | 14 days post inoculation with *S. meliloti* 1021 |
| SRR5740869 | MtNod14dpi_3 | Nodules | 14 days post inoculation with *S. meliloti* 1021 |
| SRR5740860 | MtNod14dpi_12h_1 | Nodules | 14 days post inoculation with *S. meliloti* 1021, 12h nitrogen treatment |
| SRR5740871 | MtNod14dpi_12h_2 | Nodules | 14 days post inoculation with *S. meliloti* 1021, 12h nitrogen treatment |
| SRR5740874 | MtNod14dpi_12h_3 | Nodules | 14 days post inoculation with *S. meliloti* 1021, 12h nitrogen treatment |
| SRR5740873 | MtNod14dpi_48h_1 | Nodules | 14 days post inoculation with *S. meliloti* 1021, 48h nitrogen treatment |
| SRR5740872 | MtNod14dpi_48h_2 | Nodules | 14 days post inoculation with *S. meliloti* 1021, 48h nitrogen treatment |
| SRR5740876 | MtNod14dpi_48h_3 | Nodules | 14 days post inoculation with *S. meliloti* 1021, 48h nitrogen treatment |
| SRR5740867 | Mt4wkNod_1 | Nodules | 4 weeks post inoculation with *S. meliloti* 1021 |
| SRR5740865 | Mt4wkNod_2 | Nodules | 4 weeks post inoculation with *S. meliloti* 1021 |
| SRR5740863 | Mt4wkNod_3 | Nodules | 4 weeks post inoculation with *S. meliloti* 1021 |

**Supplemental table 3.**
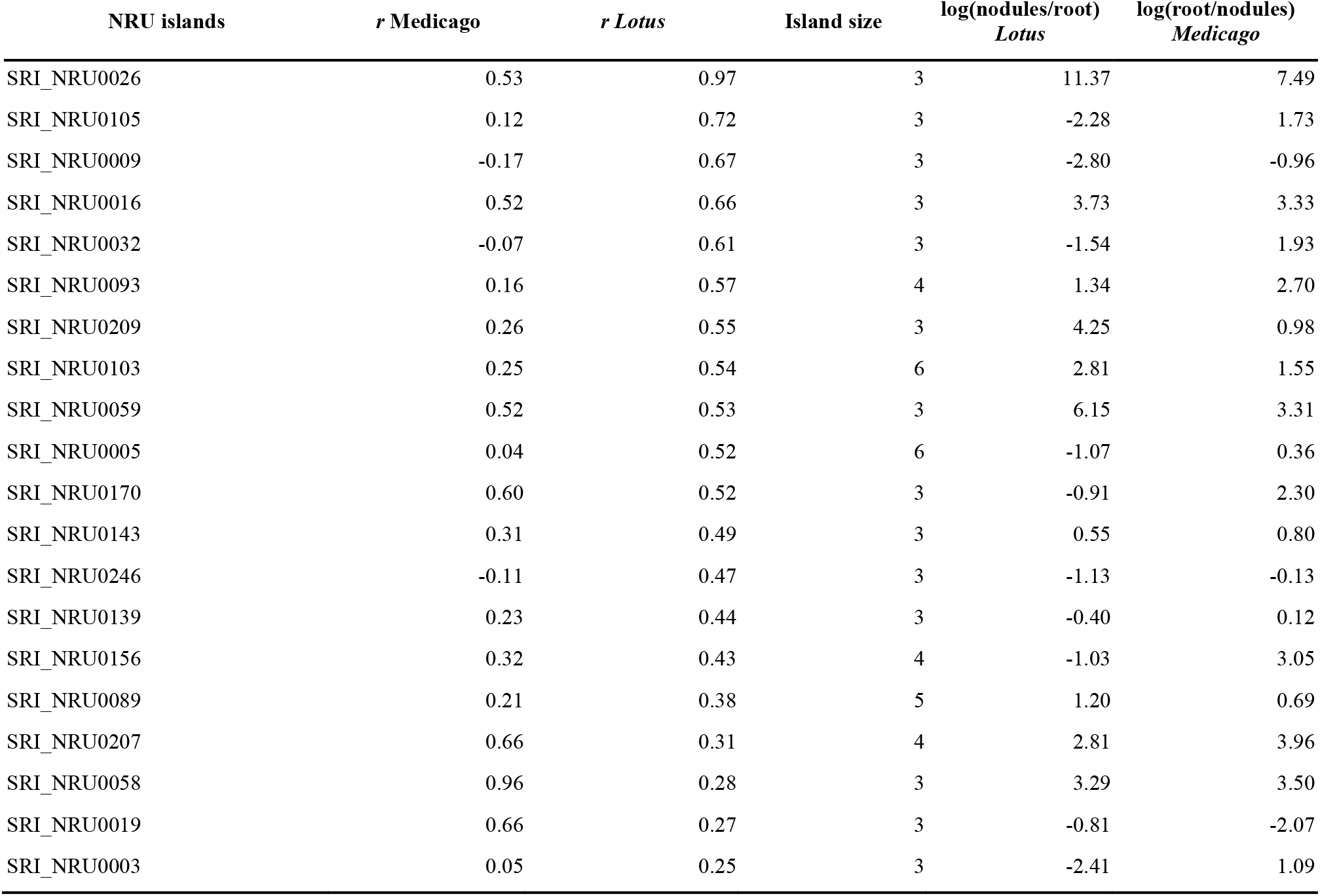
Top correlated NRU islands. The table is sorted by the *Lotus* correlation coefficient (*r*). *r*: average Pearson correlation coefficient for pairwise comparisons of gene expression profiles. Island size: number of genes within each island.

| NRU islands | *r* Medicago | *r* Lotus | Island size | log(nodules/root)<br>*Lotus* | log(root/nodules)<br>*Medicago* |
| --- | --- | --- | --- | --- | --- |
| SRI_NRU0026 | 0.53 | 0.97 | 3 | 11.37 | 7.49 |
| SRI_NRU0105 | 0.12 | 0.72 | 3 | -2.28 | 1.73 |
| SRI_NRU0009 | -0.17 | 0.67 | 3 | -2.80 | -0.96 |
| SRI_NRU0016 | 0.52 | 0.66 | 3 | 3.73 | 3.33 |
| SRI_NRU0032 | -0.07 | 0.61 | 3 | -1.54 | 1.93 |
| SRI_NRU0093 | 0.16 | 0.57 | 4 | 1.34 | 2.70 |
| SRI_NRU0209 | 0.26 | 0.55 | 3 | 4.25 | 0.98 |
| SRI_NRU0103 | 0.25 | 0.54 | 6 | 2.81 | 1.55 |
| SRI_NRU0059 | 0.52 | 0.53 | 3 | 6.15 | 3.31 |
| SRI_NRU0005 | 0.04 | 0.52 | 6 | -1.07 | 0.36 |
| SRI_NRU0170 | 0.60 | 0.52 | 3 | -0.91 | 2.30 |
| SRI_NRU0143 | 0.31 | 0.49 | 3 | 0.55 | 0.80 |
| SRI_NRU0246 | -0.11 | 0.47 | 3 | -1.13 | -0.13 |
| SRI_NRU0139 | 0.23 | 0.44 | 3 | -0.40 | 0.12 |
| SRI_NRU0156 | 0.32 | 0.43 | 4 | -1.03 | 3.05 |
| SRI_NRU0089 | 0.21 | 0.38 | 5 | 1.20 | 0.69 |
| SRI_NRU0207 | 0.66 | 0.31 | 4 | 2.81 | 3.96 |
| SRI_NRU0058 | 0.96 | 0.28 | 3 | 3.29 | 3.50 |
| SRI_NRU0019 | 0.66 | 0.27 | 3 | -0.81 | -2.07 |
| SRI_NRU0003 | 0.05 | 0.25 | 3 | -2.41 | 1.09 |

**Supplemental table 4.** Top correlated NRN islands. The table is sorted by the *Lotus* correlation coefficient (*r*). *r*: average Pearson correlation coefficient for pairwise comparisons of gene expression profiles. Island size: number of genes within each island.

| NRN islands | *r* *Medicago* | *r* *Lotus* | Island size | log(nodules/root)<br>*Lotus* | log(root/nodules)<br>*Medicago* |
| --- | --- | --- | --- | --- | --- |
| SRI_NRN0005 | 0.19 | 0.62 | 3 | -0.65 | 0.09 |
| SRI_NRN0060 | 0.14 | 0.61 | 4 | -0.96 | -0.74 |
| SRI_NRN0021 | -0.18 | 0.49 | 3 | -0.73 | 0.90 |
| SRI_NRN0034 | 0.04 | 0.46 | 3 | 0.16 | -0.44 |
| SRI_NRN0028 | 0.18 | 0.33 | 5 | 0.20 | 0.66 |
| SRI_NRN0068 | 0.06 | 0.31 | 5 | -0.24 | -0.12 |
| SRI_NRN0074 | -0.31 | 0.26 | 3 | -1.49 | -0.40 |
| SRI_NRN0044 | 0.19 | 0.24 | 3 | -0.46 | 0.26 |
| SRI_NRN0013 | -0.04 | 0.23 | 8 | 0.34 | -0.06 |
| SRI_NRN0075 | -0.03 | 0.23 | 3 | 0.53 | 0.10 |
| SRI_NRN0018 | 0.04 | 0.21 | 7 | 0.19 | -0.15 |
| SRI_NRN0014 | 0.02 | 0.20 | 3 | 0.74 | 4.06 |
| SRI_NRN0016 | 0.07 | 0.15 | 4 | -1.13 | -0.66 |
| SRI_NRN0035 | 0.19 | 0.14 | 4 | -0.22 | 0.14 |
| SRI_NRN0051 | 0.06 | 0.12 | 5 | 0.09 | -0.03 |
| SRI_NRN0038 | 0.00 | 0.10 | 7 | -0.16 | 0.30 |
| SRI_NRN0030 | 0.03 | 0.09 | 3 | 0.51 | 0.13 |
| SRI_NRN0056 | -0.23 | 0.08 | 4 | 0.22 | 0.08 |
| SRI_NRN0080 | -0.14 | 0.08 | 4 | -0.16 | -0.28 |
| SRI_NRN0079 | 0.07 | 0.07 | 4 | 0.25 | -0.08 |

**Supplemental table 5.** Top correlated NRD islands. The table is sorted by the *Lotus* correlation coefficient (*r*). *r*: average Pearson correlation coefficient for pairwise comparisons of gene expression profiles. Island size: number of genes within each island.

| NRD islands | *r* Medicago | *r* Lotus | Island size | log(nodules/root)<br>*Lotus* | log(root/nodules)<br>*Medicago* |
| --- | --- | --- | --- | --- | --- |
| SRI_NRD0027 | 0.43 | 0.88 | 5 | -4.36 | -1.63 |
| SRI_NRD0080 | 0.73 | 0.85 | 3 | -3.81 | -3.36 |
| SRI_NRD0087 | 0.71 | 0.74 | 3 | -4.08 | -3.30 |
| SRI_NRD0005 | -0.15 | 0.66 | 3 | -3.11 | -1.60 |
| SRI_NRD0017 | 0.71 | 0.64 | 4 | -2.75 | -2.91 |
| SRI_NRD0031 | -0.02 | 0.56 | 4 | -3.45 | -2.92 |
| SRI_NRD0047 | 0.21 | 0.49 | 3 | -2.70 | -0.49 |
| SRI_NRD0046 | 0.85 | 0.47 | 3 | -3.68 | -4.92 |
| SRI_NRD0045 | 0.82 | 0.37 | 3 | -4.01 | -3.62 |
| SRI_NRD0019 | 0.01 | 0.36 | 3 | -0.96 | -1.55 |
| SRI_NRD0044 | 0.22 | 0.36 | 5 | -1.47 | -2.56 |
| SRI_NRD0034 | -0.15 | 0.34 | 4 | -0.84 | -0.63 |
| SRI_NRD0043 | 0.77 | 0.34 | 3 | 2.30 | -1.35 |
| SRI_NRD0042 | 0.47 | 0.33 | 5 | -2.16 | -1.15 |
| SRI_NRD0072 | 0.49 | 0.28 | 4 | -0.86 | -3.09 |
| SRI_NRD0007 | 0.35 | 0.26 | 6 | -1.58 | -1.83 |
| SRI_NRD0006 | 0.67 | 0.20 | 3 | 1.55 | -2.10 |
| SRI_NRD0056 | -0.05 | 0.20 | 4 | -2.78 | -1.73 |
| SRI_NRD0024 | 0.55 | 0.17 | 4 | 2.99 | -2.93 |
| SRI_NRD0054 | 0.63 | 0.16 | 3 | -0.94 | -1.65 |

**Supplemental table 6.** Manually curated genes.

| gene ID | chromosome | start | end | strand |
| --- | --- | --- | --- | --- |
| LotjaGi2g1v0394950 | chr2 | 89244457 | 89246244 | - |
| LotjaGi2g1v0440600 | chr2 | 93688296 | 93690318 | + |
| LotjaGi3g1v0414350 | chr3 | 81961144 | 81962772 | - |
| LotjaGi3g1v0449330 | chr3 | 86285726 | 86285971 | + |
| LotjaGi3g1v0449360 | chr3 | 86294832 | 86295050 | + |
| LotjaGi4g1v0376650 | chr4 | 75571048 | 75571728 | + |
| LotjaGi5g1v0359230 | chr5 | 67390287 | 67391829 | + |
| LotjaGi5g1v0359260 | chr5 | 67393483 | 67394919 | + |
| LotjaGi5g1v0359300 | chr5 | 67400904 | 67402147 | + |
| LotjaGi5g1v0359350 | chr5 | 67402703 | 67404660 | - |

## References

1. Parniske, M. 2008, Arbuscular mycorrhiza: the mother of plant root endosymbioses. Nat. Rev. Microbiol., 6, 763–75.

2. Roy, S., Liu, W., Nandety, R. S., et al. 2020, Celebrating 20 Years of Genetic Discoveries in Legume Nodulation and Symbiotic Nitrogen Fixation. Plant Cell, 32, 15–41.

3. Ito, M., Miyamoto, J., Mori, Y., et al. 2000, Genome and Chromosome Dimensions of Lotus japonicus. J Plant Res, 113, 435–42.

4. Handberg, K., and Stougaard, J. 1992, Lotus japonicus, an autogamous, diploid legume species for classical and molecular genetics. Plant J., 2, 487–96.

5. Schauser, L., Roussis, A., Stiller, J., and Stougaard, J. 1999, A plant regulator controlling development of symbiotic root nodules. Nature, 402, 191–5.

6. Radutoiu, S., Madsen, L. H., Madsen, E. B., et al. 2003, Plant recognition of symbiotic bacteria requires two LysM receptor-like kinases. Nature, 425, 585–92.

7. Kawaharada, Y., Kelly, S., Nielsen, M. W., et al. 2015, Receptor-mediated exopolysaccharide perception controls bacterial infection. Nature, 523, 308–12.

8. Bruneau, A., Doyle, J. J., Herendeen, P., et al. 2013, Legume phylogeny and classification in the 21st century: Progress, prospects and lessons for other species–rich clades. TAXON, 62, 217–48.

9. Doyle, J. J. 2011, Phylogenetic Perspectives on the Origins of Nodulation. http://dx.doi.org/10.1094/MPMI-05-11-0114, 24, 1289–95.

10. Shah, N., Wakabayashi, T., Kawamura, Y., et al. 2020, Extreme genetic signatures of local adaptation during Lotus japonicus colonization of Japan. Nat Commun, 11, 253–15.

11. Sandal, N., Petersen, T. R., Murray, J., et al. 2006, Genetics of symbiosis in Lotus japonicus: recombinant inbred lines, comparative genetic maps, and map position of 35 symbiotic loci. Mol. Plant Microbe Interact., 19, 80–91.

12. Shah, N., Hirakawa, H., Kusakabe, S., et al. 2016, High-resolution genetic maps of Lotus japonicus and L. burttii based on re-sequencing of recombinant inbred lines. DNA Res., 23, 487–94.

13. Perry, J. A., Wang, T. L., Welham, T. J., et al. 2003, A TILLING Reverse Genetics Tool and a Web-Accessible Collection of Mutants of the Legume Lotus japonicus. Plant Physiol., 131, 866–71.

14. Małolepszy, A., Mun, T., Sandal, N., et al. 2016, The LORE1 insertion mutant resource. Plant J., 88, 306–17.

15. Mun, T., Bachmann, A., Gupta, V., Stougaard, J., and Andersen, S. U. 2016, Lotus Base: An integrated information portal for the model legume Lotus japonicus. Sci Rep, 6, 39447.

16. Stougaard, J. 2014, Background and History of the Lotus japonicus Model Legume System In: The Lotus japonicus Genome. Springer, Berlin, Heidelberg, Berlin, Heidelberg, pp. 3–8.

17. Sato, S., Nakamura, Y., Kaneko, T., et al. 2008, Genome Structure of the Legume, Lotus japonicus. DNA Res., 15, 227–39.

18. Kelly, S., Mun, T., Stougaard, J., Ben, C., and Andersen, S. U. 2018, Distinct Lotus japonicus Transcriptomic Responses to a Spectrum of Bacteria Ranging From Symbiotic to Pathogenic. Front Plant Sci, 9, 1218.

19. Malolepszy, A., Kelly, S., Sørensen, K. K., et al. 2018, A plant chitinase controls cortical infection thread progression and nitrogen-fixing symbiosis. Elife, 7, e00013.

20. Zhang, H.-B., Zhao, X., Ding, X., Paterson, A. H., and Wing, R. A. 1995, Preparation of megabase-size DNA from plant nuclei. Plant J., 7, 175–84.

21. Koren, S., Walenz, B. P., Berlin, K., Miller, J. R., Bergman, N. H., and Phillippy, A. M. 2017, Canu: scalable and accurate long-read assembly via adaptive k-mer weighting and repeat separation. Genome Res., 27, 722–36.

22. Li, H., and Durbin, R. 2009, Fast and accurate short read alignment with Burrows-Wheeler transform. Bioinformatics, 25, 1754–60.

23. Li, H., Handsaker, B., Wysoker, A., et al. 2009, The Sequence Alignment/Map format and SAMtools. Bioinformatics, 25, 2078–9.

24. Lieberman-Aiden, E., van Berkum, N. L., Williams, L., et al. 2009, Comprehensive mapping of long-range interactions reveals folding principles of the human genome. Science, 326, 289–93.

25. Bickhart, D. M., Rosen, B. D., Koren, S., et al. 2017, Single-molecule sequencing and chromatin conformation capture enable de novo reference assembly of the domestic goat genome. Nat. Genet., 49, 643–50.

26. Burton, J. N., Adey, A., Patwardhan, R. P., Qiu, R., Kitzman, J. O., and Shendure, J. 2013, Chromosome-scale scaffolding of de novo genome assemblies based on chromatin interactions. Nat. Biotechnol., 31, 1119–25.

27. Gremme, G., Brendel, V., Sparks, M. E., and Kurtz, S. 2005, Engineering a software tool for gene structure prediction in higher organisms. Information and Software Technology, 47, 965–78.

28. Kim, D., Paggi, J. M., Park, C., Bennett, C., and Salzberg, S. L. 2019, Graph-based genome alignment and genotyping with HISAT2 and HISAT-genotype. Nat. Biotechnol., 37, 907–15.

29. Pertea, M., Pertea, G. M., Antonescu, C. M., Chang, T.-C., Mendell, J. T., and Salzberg, S. L. 2015, StringTie enables improved reconstruction of a transcriptome from RNA-seq reads. Nat. Biotechnol., 33, 290–5.

30. Altschul, S. F., Gish, W., Miller, W., Myers, E. W., and Lipman, D. J. 1990, Basic local alignment search tool. J. Mol. Biol., 215, 403–10.

31. Eddy, S. R. 2011, Accelerated Profile HMM Searches. Pearson, W. R., (ed.). PLoS Comput Biol, 7, e1002195.

32. Tomato Genome Consortium. 2012, The tomato genome sequence provides insights into fleshy fruit evolution. Nature, 485, 635–41.

33. Smit A, Hubley R, Green P. RepeatMasker Open-4.0. 2013–2015. www.repeatmasker.org.

34. Lowe, T. M., and Eddy, S. R. 1997, tRNAscan-SE: a program for improved detection of transfer RNA genes in genomic sequence. Nucleic Acids Res., 25, 955–64.

35. Lagesen, K., Hallin, P., Rødland, E. A., Staerfeldt, H.-H., Rognes, T., and Ussery, D. W. 2007, RNAmmer: consistent and rapid annotation of ribosomal RNA genes. Nucleic Acids Res., 35, 3100–8.

36. Nawrocki, E. P., and Eddy, S. R. 2013, Infernal 1.1: 100-fold faster RNA homology searches. Bioinformatics, 29, 2933–5.

37. Patro, R., Duggal, G., Love, M. I., Irizarry, R. A., and Kingsford, C. 2017, Salmon provides fast and bias-aware quantification of transcript expression. Nat. Methods, 14, 417–9.

38. Love, M. I., Huber, W., and Anders, S. 2014, Moderated estimation of fold change and dispersion for RNA-seq data with DESeq2. Genome Biol., 15, 550–21.

39. Pecrix, Y., Staton, S. E., Sallet, E., et al. 2018, Whole-genome landscape of Medicago truncatula symbiotic genes. Nat Plants, 356, eaad4501.

40. Dobin, A., Davis, C. A., Schlesinger, F., et al. 2013, STAR: ultrafast universal RNA-seq aligner. Bioinformatics, 29, 15–21.

41. Anders, S., Pyl, P. T., and Huber, W. 2015, HTSeq--a Python framework to work with high-throughput sequencing data. Bioinformatics, 31, 166–9.

42. Hayashi, M., Miyahara, A., Sato, S., et al. 2001, Construction of a Genetic Linkage Map of the Model Legume Lotus japonicus Using an Intraspecific F 2 Population. DNA Res., 8, 301–10.

43. Ohmido, N., Ishimaru, A., Kato, S., Sato, S., Tabata, S., and Fukui, K. 2010, Integration of cytogenetic and genetic linkage maps of Lotus japonicus, a model plant for legumes. Chromosome Res., 18, 287–99.

44. Simão, F. A., Waterhouse, R. M., Ioannidis, P., Kriventseva, E. V., and Zdobnov, E. M. 2015, BUSCO: assessing genome assembly and annotation completeness with single-copy orthologs. Bioinformatics, 31, 3210–2.

45. Munch, D., Gupta, V., Bachmann, A., et al. 2018, The Brassicaceae Family Displays Divergent, Shoot-Skewed NLR Resistance Gene Expression. Plant Physiol., 176, 1598–609.

46. Handa, Y., Nishide, H., Takeda, N., Suzuki, Y., Kawaguchi, M., and Saito, K. 2015, RNA-seq Transcriptional Profiling of an Arbuscular Mycorrhiza Provides Insights into Regulated and Coordinated Gene Expression in Lotus japonicus and Rhizophagus irregularis. Plant Cell Physiol., 56, 1490–511.

47. Schmutz, J., Cannon, S. B., Schlueter, J., et al. 2010, Genome sequence of the palaeopolyploid soybean. Nature, 463, 178–83.

48. Høgslund, N., Radutoiu, S., Krusell, L., et al. 2009, Dissection of symbiosis and organ development by integrated transcriptome analysis of lotus japonicus mutant and wild-type plants. Provart, N. J., (ed.). PLoS ONE, 4, e6556.

49. Verdier, J., Torres-Jerez, I., Wang, M., et al. 2013, Establishment of the Lotus japonicus Gene Expression Atlas (LjGEA) and its use to explore legume seed maturation. Plant J., 74, 351–62.

50. Kereszt, A., Mergaert, P., Montiel, J., Endre, G., and Kondorosi, É. 2018, Impact of Plant Peptides on Symbiotic Nodule Development and Functioning. Front Plant Sci, 9, 1026.

51. Goodstein, D. M., Shu, S., Howson, R., et al. 2012, Phytozome: a comparative platform for green plant genomics. Nucleic Acids Res., 40, D1178–86.

52. Dash, S., Campbell, J. D., Cannon, E. K. S., et al. 2016, Legume information system (LegumeInfo.org): a key component of a set of federated data resources for the legume family. Nucleic Acids Res., 44, D1181–8.

53. Lyons, E., and Freeling, M. 2008, How to usefully compare homologous plant genes and chromosomes as DNA sequences. Plant J., 53, 661–73.

54. Griesmann, M., Chang, Y., Liu, X., et al. 2018, Phylogenomics reveals multiple losses of nitrogen-fixing root nodule symbiosis. Science, 361, eaat1743.

## Supplemental references

Handa, Y., Nishide, H., Takeda, N., Suzuki, Y., Kawaguchi, M., and Saito, K. (2015). RNA-seq Transcriptional Profiling of an Arbuscular Mycorrhiza Provides Insights into Regulated and Coordinated Gene Expression in Lotus japonicus and Rhizophagus irregularis. Plant Cell Physiol. 56, 1490–1511.

Kelly, S., Mun, T., Stougaard, J., Ben, C., and Andersen, S.U. (2018). Distinct Lotus japonicus Transcriptomic Responses to a Spectrum of Bacteria Ranging From Symbiotic to Pathogenic. Front Plant Sci 9, 1218.

Munch, D., Gupta, V., Bachmann, A., Busch, W., Kelly, S., Mun, T., and Andersen, S.U. (2018). The Brassicaceae Family Displays Divergent, Shoot-Skewed NLR Resistance Gene Expression. Plant Physiol. 176, 15981609.

