## Supplemental Data 1 for "Insights into the evolution of symbiosis gene copy number and distribution from a chromosome-scale *Lotus japonicus* Gifu genome sequence": short_summary_busco_1.2_12072019.txt

# BUSCO version is: 3.0.2 
# The lineage dataset is: embryophyta_odb9 (Creation date: 2016-02-13, number of species: 30, number of BUSCOs: 1440)
# To reproduce this run: python /home/pgsb/nadia.kamal/tools/busco/scripts/run_BUSCO.py -i LjGifu1.2_aa.fasta -o busco_1.2_12072019 -l /home/pgsb/nadia.kamal/tools/busco/embryophyta_odb9/ -m proteins -c 1
#
# Summarized benchmarking in BUSCO notation for file LjGifu1.2_aa.fasta
# BUSCO was run in mode: proteins

	C:94.2%[S:61.1%,D:33.1%],F:0.8%,M:5.0%,n:1440

	1357	Complete BUSCOs (C)
	880	Complete and single-copy BUSCOs (S)
	477	Complete and duplicated BUSCOs (D)
	12	Fragmented BUSCOs (F)
	71	Missing BUSCOs (M)
	1440	Total BUSCO groups searched
