## Supplemental Data 1 for "Insights into the evolution of symbiosis gene copy number and distribution from a chromosome-scale *Lotus japonicus* Gifu genome sequence": README.txt

Note that the version number 1.1 on the LjGifu1.1_pseudomol.fa genome reference file refers to the assembly version, while the 1.2 version number on the remaining files refer to the annotation version with reference to the version 1.1 assembly.
